# Loss of SRF-driven contractility, not lymphatic smooth muscle cell loss, is the primary driver of age-associated collecting vessel decline

**DOI:** 10.64898/2026.11.19.767563

**Authors:** Michael.M. Orlich, Amanda Norrén, Hans Schoofs, Malavika Desai, Ceren Saygi, Soumiya Pal, Yixiao Liu, Scott D. Zawieja, Taija Mäkinen, Konstantin Gängel

## Abstract

The lymphatic vasculature returns interstitial fluid to the blood circulation, a function that depends on rhythmic contraction of the smooth muscle cells (SMCs) surrounding collecting vessels. Lymph transport declines with age, a decline attributed largely to loss of SMCs from collecting vessels. We show that the principal change is not their loss but the loss of their contractile phenotype. In mouse ear collecting vessels, contractile protein expression declined with age and the vessels dilated, yet the majority of SMCs remained on the vessel wall. The decline extended to serum response factor (SRF), the transcription factor driving the smooth muscle contractile gene programme, and was mirrored by a loss of SRF-dependent contractile transcripts in published single-cell data from aged lymphatic muscle cells. Inducible deletion of *Srf* in mural cells reproduced this state, abolishing the contractile programme while leaving the cells in place and thereby excluding impaired SMC recruitment as the main mechanism. The resulting vessels became severely dilated and misshapen, whereas lymphatic capillaries were unaffected; endothelial identity, valve formation and lymphatic endothelial cell (LEC) number were all preserved, with enlargement instead reflecting increased LEC size. Intravital imaging showed loss of both rhythmic contraction and the contractile movement of individual SMCs, and Evans blue drainage to the inguinal lymph node was correspondingly reduced. Deletion of *Srf* after the contractile phenotype was established recapitulated this loss, showing that SRF is required continuously to maintain lymphatic SMC contractility and providing a genetic model of age-associated lymphatic dysfunction.

**Statements and Declarations:** The authors declare that they have no conflict of interest.

## Introduction

In contrast to the blood vasculature, the lymphatic vasculature is organized in a blind-ended fashion. Lymphatic capillaries take up interstitial fluid, thereafter termed lymph, and transport it to pre-collecting and collecting vessels. These collecting vessels contain valves that enable unidirectional lymph transport towards the lymph nodes. From there, lymph is transported to the sole anatomical connection between the two vascular systems, the subclavian vein^1-3^. While collecting vessels can be innervated to support the regulation of lymph flow^4^, they can also function as autonomous pumps, generating spontaneous and rhythmic contractions^5^. This propulsive activity is generated by lymphatic smooth muscle cells (lSMCs). Similar to vascular smooth muscle cells (vSMCs) and pericytes in the blood vasculature, lSMCs belong to the group of mural cells. Analogous to mural cell recruitment in the blood vasculature, lymphatic endothelial cells of collecting vessels recruit lSMCs through the secretion of platelet-derived growth factor B (PDGFB)^6^.

Collecting vessel function declines with age, a decline that has been attributed in large part to the loss of lSMCs from the vessel wall. Its consequences are of direct clinical relevance, because dysfunctional lymphatic drainage leads to the accumulation of lymph in tissues and ultimately results in lymphedema. Although lymphedema is a multifactorial disease^7^, accumulating evidence suggests the involvement of lSMCs^8^. For example, gain-of-function mutations in smooth muscle ATP-sensitive potassium (KATP) channels cause Cantú syndrome, a multisystem disorder frequently accompanied by lymphedema, in which lSMC contraction is impaired^9, 10^. Furthermore, lSMCs from patients with late-stage chronic lymphedema show reduced levels of the smooth muscle myosin heavy chain isoforms SM1 and SM2, indicating impaired contractile function^11^. Reduced lymphatic pumping in aged animals has been documented in both mice and rats^12-16^, and single-cell RNA sequencing of peripheral collecting lymphatic vessels showed reduced expression of genes encoding the contractile apparatus in lSMCs from aged mice^15^. Besides the increased risk of lymphedema, reduced lymphatic pumping also impairs antigen transport and is associated with immunosuppression ^13^.

Current treatments for lymphedema remain largely supportive and are often insufficient. Invasive surgeries such as the lymphatic-venous anastomosis are often the last option for patients with severe lymphedema, and carry the risk of blood refluxing into the lymphatic vessel^11^. The transcriptional programmes that specify and maintain lymphatic endothelial identity, such as those governed by PROX1 and FOXC2, are well defined. By contrast, how the contractile phenotype of the surrounding muscle layer is transcriptionally controlled, and why it deteriorates during aging and in chronic lymphedema, has remained largely unexplored^1, 17-23^. Genetic models have shown that preventing lSMC recruitment, through deletion of endothelial *Pdgfb*, produces dilated collecting vessels that fail to contract^6^. Whether the age-associated decline of collecting vessel function likewise reflects the disappearance of lSMCs, or instead a loss of the contractile phenotype of lSMCs that remain on the vessel wall, has not been resolved, not least because αSMA has commonly been used both to identify these cells and to report their contractile state. Defining the transcriptional programme that sustains lSMC contractility is therefore a prerequisite for distinguishing these possibilities and for any attempt to restore lymphatic pumping therapeutically.

The smooth muscle contractile gene programme in the blood vasculature is governed by the transcription factor serum response factor (SRF), which binds CArG box elements in the promoters of genes encoding the contractile apparatus together with its co-factor myocardin^24-29^. We previously showed that SRF controls the contractile phenotype, and thereby the vascular tone, of mural cells in the blood vasculature^30^. Postnatal mural cell-specific deletion of *Srf* caused a loss of vSMC contractility and, during the remodeling phase of the murine retinal vasculature, triggered the formation of arteriovenous shunts, which develop in response to a lack of myogenic activity and relieve the resulting rise in pressure. Because lSMCs deploy a similar contractile machinery^31^, we reasoned that SRF was a strong candidate for the transcriptional control of the lSMC contractile phenotype, and that removing it would isolate a single variable: whether loss of mural cell contractility alone is sufficient to compromise collecting vessel structure and lymph transport.

Here we show that the dominant change in aged collecting vessels is not the major disappearance of lSMCs but the loss of their contractile phenotype, and we identify SRF as the transcriptional regulator that sustains it. lSMCs lacking SRF remained associated with the vessel wall despite losing their contractile machinery, and this alone was sufficient to distort collecting vessel architecture and to abolish the rhythmic contractions that drive lymph transport. The requirement for SRF persists after the contractile phenotype is established, so that deleting *Srf* in adult animals reproduces the lSMC phenotype of aged mice and provides a genetic model in which age-associated lymphatic decline can be studied.

## Materials and methods

### Ethical Statement

Animal housing and the experiments performed were in accordance with the Swedish and German legislation and were approved by the local animal ethics committees before experimentation. The protocols used in this study were approved by the Uppsala Committee (local approval numbers: 5.8.18-03029-2020, 5.8.18-06383/2020, and 5.8.18-03362/2021) and the Behörde für Justiz und Verbraucherschutz der Freien und Hansestadt Hamburg, Germany (local approval number: N033/2025). All the animal procedures have been executed conform to the Directive 2010/63/EU of the European Parliament on the protection of animals used for scientific purposes. All efforts were made to minimize animal suffering. Both male and female mice (in equal proportion) were used in this study if not stated otherwise.

### Animal housing

Mice were held in individually ventilated cages in groups of 2 to 5 animals per cage with a 12-hour day/night cycle (6 am to 6 pm), a relative humidity of 60%, and a temperature of 21 °C. Animals had unrestricted access to food (pellets, composition: crude protein, 19.3%; crude fat, 3.3%; crude fiber, 4.4%; crude ash, 6.0%; calcium, 1.0%; phosphorus, 0.7%, low in phytoestrogens) and water ad libitum. New litters were weaned 3 weeks after birth and separated into new cages.

### Mouse Lines and Inducible Genetic experiments

Tamoxifen-inducible mural cell-specific *Srf* knockout (*Srf*^*iMCKO*^) mice were generated by breeding *Pdgfrb(BAC)-CreER*^*T2*^ (Tg(Pdgfrb-cre)1Rha) mice with *Srf*^*fl/fl*^ *(Srf*^*tm2*.*1Nor*^) mice. Similar, adult vSMC specific *Srf* knockout (*Srf*^*ivSMCKO*^) were generated by breeding *Itga8-CreER*^*T2*^ (*Itga8*^tm1.1(cre/ERT2)Jomm^) with *Srf*^*fl/fl*^ mice. All mice were bred on a *C57BL/6J* genetic background (backcrossed at least 4 generations). Both male and female mice were used for analysis, and no difference in the phenotype between the sexes was observed. For postnatal analysis. The day of birth was considered postnatal day (P) 0. Cre-mediated recombination was induced with 150 µg tamoxifen (Sigma T5648) dissolved in peanut oil (Sigma P2144) by direct feeding at postnatal days P1 to P3. *Srf*^*iMCKO*^ (*Pdgfrb(BAC)*-*CreER*^*T2*^; *Srf*^*fl/fl*^) mice were compared with Cre-positive *Srf*-heterozygous control (*Pdgfrb(BAC)-CreER*^*T2*^; *Srf*^*fl/wt*^) mice to exclude potential Cre-induced toxicity as the cause of the observed phenotype^32^. For some experiments, Cre-negative control (*Srf*^*fl/fl*^ or *Srf*^*fl/wt*^) mice were used as additional controls. Furthermore, for analyses of lSMC morphology we additionally crossed the Rosa^mTmG^ (Gt(ROSA)26Sor^tm4(ACTB-tdTomato,-EGFP)Luo^) reporter into the described line and compared only Cre-positive *Srf*-heterozygous control to Cre-positive *Srf*-homozygous knockouts. For adult *Srf-Itga8CreER*^*T2*^ inductions, 1 mg of tamoxifen (dissolved in peanut oil) was injected intraperitoneally on five consecutive days. For the analysis of aged ear skins shown in Figure 1, C57BL/6J wild-type mice from our in-house breeding facility were used.

**Figure 1.**
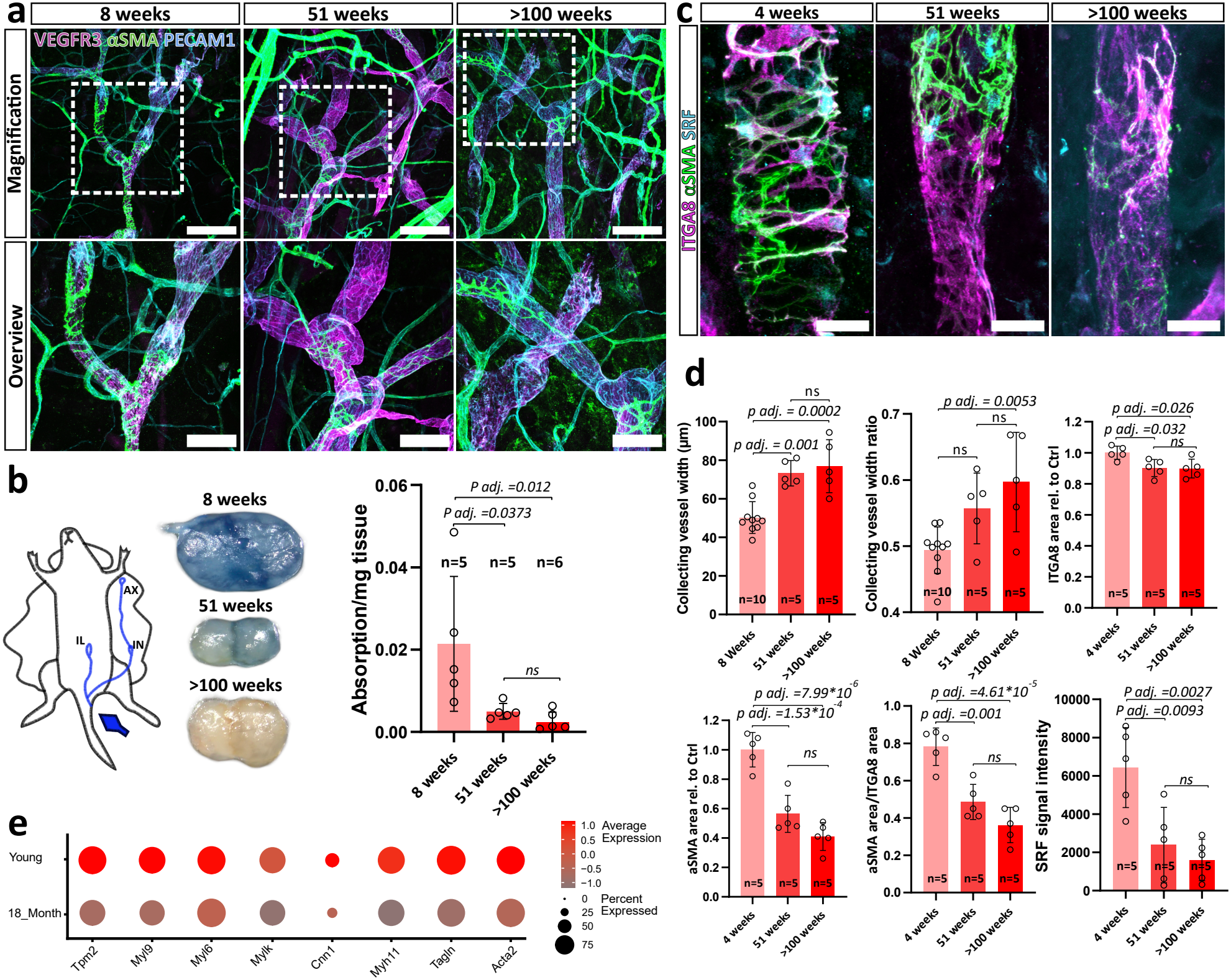
Morphometric changes in mouse ear collecting vessels and reduced expression of SRF-regulated contractile proteins in aged lymphatic SMCs. **(a)** High-resolution confocal images of wild-type lymphatic collecting vessels (VEGFR3, magenta; PECAM1, cyan) from mice of different ages, showing expression of the contractile protein αSMA (green). Note the increased vessel diameter in older mice. Scale bars, 200 µm (overview) and 100 µm (magnified view). **(b)** Functional lymphatic drainage assessment via subcutaneous tail injection of Evans blue dye (schematic, upper left) and Evans blue content in inguinal lymph nodes (right graph, number of animals (n)= 5 (8/51 weeks), and n=6 (>100 weeks)). Inguinal lymph nodes were collected 5 min post-injection. **(c)** Confocal images of lymphatic collecting vessels from 4-, 51-, and >100-week-old mice showing αSMA (magenta) and SRF (green). Note the reduced SRF signal in aged lymphatic SMCs. Scale bar, 25 µm. **(d)** Morphometric analysis of collecting vessel diameter and vessel width ratio (upper graphs; quantified from a, n= 10 (8weeks) and 5 (51 weeks and >100 weeks), as well as αSMA/ITGA8-positive area (n=5, each group) and SRF signal intensity (n=5, each group) in lymphatic SMCs (lower graphs; quantified from c) in wild-type ear collecting vessels of different ages. **(e)** Dot plot showing expression of SRF-regulated contractile genes in young versus aged lymphatic SMCs based on the published single-cell RNA sequencing dataset of Lei and Ruscic et al. (2025). Error bars represent s.d. of the mean. Statistical significance in **b** and **d** was determined using the Shapiro-Wilk test for normality and a one-way ANOVA multiple comparisons test with Tukey’s-correction (3 comparisons). P-values are indicated.

### Tissue preparation and Immunohistology

Ear tissues were dissected from postnatal mice and separated into dorsal and ventral sides. Only the dorsal side was further used. Excessive cartilage was removed and tissues were fixed in 4% paraformaldehyde (Sigma; No. P6148) dissolved in PBS for 1 hour at room temperature. Afterwards, ear tissues were washed in PBS followed by blocking and permeabilization in permblock solution: PBS (Gibco 14190-094) supplemented with 1% BSA (Sigma; No. A4378) and 0.3% Triton X-100 (Sigma; No. T8787) for 2 hours at room temperature. Primary antibodies were diluted in permblock buffer supplemented with 2% donkey serum (Sigma; No. S30-M) and incubated overnight at 4 °C. The next day, the primary antibody solution was removed and the ear samples were washed 3× for at least 20 minutes using PBS. Samples were then incubated with fluorescence-coupled secondary antibodies in permblock buffer supplemented with 2% donkey serum for 2 hours at RT protected from light. After incubation, ear tissues were washed 3× for at least 20 minutes with PBS. Finally, samples were mounted on a microscope slide using Fluoromount-G (Southern Biotech; No. 0100-01) mounting medium. Microscopy was performed on a Leica SP8 confocal and a Zeiss LSM 980 Airy Scan 2 (DFG code: INST 152/876-1 FUGG) setup. Image analysis was performed using Fiji/ImageJ (version 1.54f)^33^.

### Evans blue lymphatic drainage assay

For quantification of inguinal lymphatic drainage, approx. 30 µl of Evans blue (2%; Sigma-Aldrich) was injected subcutaneously at the base of the mouse tail skin. After 5 min, the mice were sacrificed and both inguinal lymph nodes were subsequently dissected and pooled. The weight of the lymph nodes was noted. Afterwards, lymph nodes were incubated in 100 µl of formamide for 24 h at 60 °C to isolate the Evans blue dye from the tissue. Afterwards, absorption was measured photometrically at 620 nm and normalized to the tissue weight.

### In vivo proliferation assay

In vivo lSMC proliferation was assessed as previously described by Wang et al. (2017)^6^. 5-Ethynyl-2⍰-deoxyuridine (EdU; 100 µg/mouse) was administered intraperitoneally at P14, P16, and P18. Animals were sacrificed at P21, and ear skin was collected for immunofluorescence analysis. EdU incorporation was detected using the Click-iT EdU Imaging Kit according to the manufacturer’s instructions, with the incubation time of the reaction cocktail extended to 3 h.

### Intravital Microscopy

Intravital imaging was performed as previously described^34, 35^, with modifications. In short, mice were anesthetized once using ketamine (66.7 mg/kg) and xylazine (11.7 mg/kg) via intraperitoneally injection route. Anesthetized animals were placed on a heating pad and the eyes were moisturized with an eye cream (Bepanthen; Bayer Vital GmbH, Germany). Afterward, 2 µl of fluorescein isothiocyanate–labeled dextran (500 kDa, 10 mg/mL; FD500; TdB Labs, Sweden) was injected subcutaneously to the tip of the ear using a 10 µl Hamilton syringe. Next, the animal was placed under an upright multiphoton microscope (LEICA SP8 DIVE), and the ventral side of the ear was fixed with double-sided tape and positioned under the objective (HC IRAPO L 25×/NA = 1.00 W motCORR; Leica). Images and image sequences were taken from the dorsal side of the ear at a resolution of 1024×1024 and a refresh time of 0.84 s per image (imaging speed, 600 Hz; bidirectional imaging; zoom, 1.0). Image analysis was performed using Fiji/ImageJ (version 1.54f) ^33^.

### Euthanasia

Mice were anaesthetized with 2-2.5% of isoflurane (except for intravital microscopy, see according chapter) and euthanized via cervical dislocation.

### Morphometric Analysis

Vessel area, capillary vessel width, and branching points were quantified from whole-ear tile scans in a blinded fashion using the REAVER GUI in MATLAB^36^. Collecting vessel width was measured manually in Fiji using the Freehand Line tool at the widest point of the vessel sinus. To determine the vessel width ratio, the narrowest point immediately distal to the sinus was measured, and the ratio of the narrowest to widest vessel diameter was calculated. On average, 10 independent collecting vessels were analyzed per animal. lSMCs were manually counted based on αSMA and Desmin staining and normalized to vessel area. Desmin coverage was determined by thresholding the Desmin-positive area and normalizing it to the VEGFR3-positive area. Similarly, αSMA coverage was calculated by thresholding the αSMA-positive area and normalizing it to the VEGFR3-positive area. Proliferating lSMCs were identified based on αSMA- or Desmin-positive staining in combination with EdU labeling and expressed as a percentage of the total number of lSMCs counted along an entire lymphatic branch. Lymphatic endothelial cell (LEC) density was determined by counting PROX1-positive cells and normalizing the cell number to the analyzed vascular area. LEC density per vessel length was calculated by dividing the number of PROX1-positive cells by the analyzed length of the lymphatic vasculature. LYVE1-positive area was quantified by thresholding and normalizing to the VEGFR3-positive area to assess lymphatic vessel maturation and identity. LEC size in collecting vessels was determined by measuring individual cell areas based on VE-cadherin staining. Lymphatic valves were identified by integrin α9-positive structures and quantified relative to the analyzed vessel length. SRF expression was quantified by measuring the mean gray value of the nuclei of integrin α8-positive lSMCs in collecting vessels and subtracting the corresponding background intensity. Vessel diameter during intravital microscopy was quantified using the Fiji Macro VasoMetrics^37^.

### Single-cell RNA-seq analysis

Publicly available single-cell RNA-seq data from murine collecting lymphatic vessels (GEO: GSE248658)^15^ were analyzed using Seurat^38^, following the analytical workflow described in the original study^15^. The six samples were individually processed and subsequently merged. The Seurat object was annotated using the author-provided cell-level metadata. Lymphatic muscle cells (LMCs) were identified based on the author-provided annotations and subjected for downstream analysis. The expression of selected genes of interest was assessed in LMCs and visualized using Seurat’s DotPlot function^38^, with cells grouped according to age group.

### Statistics

Statistical analyses were performed using the GraphPad Prism 9 software. If not otherwise stated, all data are presented as mean ± SD. Unless otherwise stated, dots represent the number of animals. When possible (n≥5), normal distribution of data was tested by using the Shapiro-Wilk test (α=0.05). For statistical comparison, unless otherwise stated, an unpaired Mann-Whitney test was performed (α=0.05, CI of 95%). For the Evans-blue perfusion test (Figure 5c), a Shapiro-Wilk test was performed to verify normality of data distribution, followed by a two-tailed Welch’s test. A *p*<0.05 was considered statistically significant. Blinding of samples was not performed, and groups were randomly assigned. No prior statistical group size determination was performed. Instead, group size was determined according to previous experience and reproducibility of the results across the independent experiments. No animals were excluded from the study. Images were chosen according to the most representative examples reflecting the typical phenotype.

### Use of artificial intelligence

ChatGPT (GPT-5.6 Luna, OpenAI) and Claude 1.32885.1 (a757f5, Anthropic) was used to improve the readability, clarity, and language of the manuscript. All AI-assisted text was reviewed and edited by the authors, who verified its accuracy and scientific content. The authors take full responsibility for the content of the manuscript.

## Acknowledgements

We thank C. Olsson, H. Leksell, J. Chmielniakova, V, Sundell, and S.L. Segerqvist for technical assistance. Intravital imaging was performed at the BioVis platform of Uppsala University and at the UMIF platform of the Medical hospital Hamburg-Eppendorf. The *Pdgfrb(BAC)-CreER*^*T2*^mouse line was kindly provided by Ralf Adams. The *Itga8-CreERT*^*T2*^ line was kindly provided by Joseph Miano and Helle Jørgensen.

## Competing interests

The authors declare no conflict of interest.

## Funding

This study was supported by the Swedish Research Council (2021-04896, K. Gaengel; 2020-0269, T. Mäkinen), the Tore Nilsons Stiftelse för Medicinsk Forskning (2020-00873, K. Gaengel), the Swedish Cancer Society (254918, K.Gaengel), the Knut and Alice Wallenberg Foundation (2020.0057, K. Gaengel and T. Mäkinen), the Gustaf Adolf Johansson’s foundation (890622-p133, M.M. Orlich), the Deutsche Forschungsgemeinschat (DFG, German Research Foundation 503902844 and 572113215, M.M. Orlich), the European Union’s Horizon 2020 research and innovation programme under the Marie Skłodowska-Curie grant agreement (no. 814316; H. Schoofs, T. Mäkinen) and NIH/NHLBI R01 - HL175083 (S.D. Zawieja).

## Data availability

A The data that support the findings of this study are available from the corresponding author upon reasonable request

## Results

### Aging reduces contractile protein expression in lymphatic SMCs that remain on collecting vessels

To determine how the lSMC compartment of collecting vessels changes with age, we analyzed lymphatic collecting vessels in the ear skin of wild-type mice at different ages and assessed the expression of contractile proteins and SRF. Ear tissues from young (4-8 weeks), middle-aged (approx. 51 weeks), and aged (>100 weeks) mice were isolated and analyzed by high-resolution confocal microscopy following immunostaining. Platelet endothelial cell adhesion molecule 1 (PECAM1) was used as a pan-endothelial marker and vascular endothelial growth factor receptor 3 (VEGFR3) as a lymphatic endothelial marker to visualize the lymphatic vasculature, while αSMA was used as a marker of the SMC contractile phenotype. Morphometric analysis revealed a significant increase in collecting vessel diameter accompanied by reduced αSMA coverage in aged mice (Figure 1a, d, upper graphs), suggesting age-dependent alterations in the contractile phenotype of lymphatic collecting vessels.

To further assess whether these morphological changes were associated with impaired lymphatic drainage, we performed an Evans blue lymphatic drainage assay. Evans blue dye was injected subcutaneously into the tail skin, from where it drains toward the inguinal lymph nodes (Figure 1b, left schematic). Five minutes after injection, the inguinal lymph nodes were isolated and Evans blue content was quantified photometrically. Consistent with the age-dependent changes observed in the ear lymphatic vasculature, Evans blue accumulation in the inguinal lymph nodes was reduced in aged mice, indicating impaired lymphatic drainage with increasing age (Figure 1b).

To investigate a potential relationship between SRF expression and the contractile phenotype of lSMCs, we next performed whole-mount immunostaining of wild-type ear tissues for the vSMC and lSMC marker integrin α8 ^39, 40^, αSMA, and SRF. This combination allowed us to identify lSMCs in collecting vessels independently of their αSMA expression, as integrin α8 expression appeared to remain relatively stable in lSMCs during aging (Figure 1d, upper graph). Analysis of these tissues revealed that SRF expression decreased with age, concomitant with a remarkable reduction of contractile protein αSMA in lymphatic SMCs (Figure 1c, d) while the integrin α8 expression only moderately dropped, suggesting a potential role for SRF in maintaining the lSMC contractile phenotype.

To further investigate this association, we re-analyzed a recently published single-cell RNA sequencing dataset by Lei and Ruscic et al. (2025)^15^, comprising lSMCs isolated from popliteal lymphatic vessels of young and aged wild-type mice. We focused on established SRF-regulated genes and observed a significant age-dependent reduction in multiple SRF target genes encoding the contractile apparatus (Figure 1e). Together, these findings indicate that aging is associated with reduced SRF expression and a concomitant reduction in the expression of contractile genes in lSMCs. We therefore hypothesized that reduced SRF activity may contribute to the age-dependent loss of the lymphatic SMC contractile phenotype. Because these aging-associated changes are correlative, we next asked whether loss of SRF is sufficient to cause this phenotype, and therefore turned to a genetic loss-of-function approach.

### Mural *Srf* deletion causes severe lymphatic collecting vessel abnormalities

To determine whether the loss of SRF is sufficient to induce structural and functional alterations of the lymphatic vasculature resembling those seen during aging and in lymphatic disease, we deleted *Srf* in mural cells. Recently, we developed a mouse model enabling efficient deletion of *Srf* in mural cells by crossing mice carrying a floxed Srf allele with *Pdgfrb-CreER*^*T2*^ mice ^30^. Lymphatic SMCs also express PDGFRβ ^6, 15, 17, 41^, and *Pdgfrb-CreER*^*T2*^ has been shown to effectively label LMCs in addition to lymphatic fibroblasts^41^. We reasoned that the same mouse line and tamoxifen-induction protocol could be used to investigate the consequences of *Srf* deletion in lSMCs.

Mural cell-specific deletion of *Srf*, hereafter referred to as *Srf*^*iMCKO*^ (inducible mural cell knockout), was induced in newborn mice by tamoxifen administration on postnatal days 1-3 (Figure 2a). We focused our analysis on the lymphatic vasculature of the ear skin at postnatal day 21, as this tissue represents a well-established model for analyzing lymphatic vessel development and morphology^15, 42-46^. At this developmental stage, the ear lymphatic vasculature is established and functional^43^, and its collecting vessels are covered by lSMCs.

**Figure 2.**
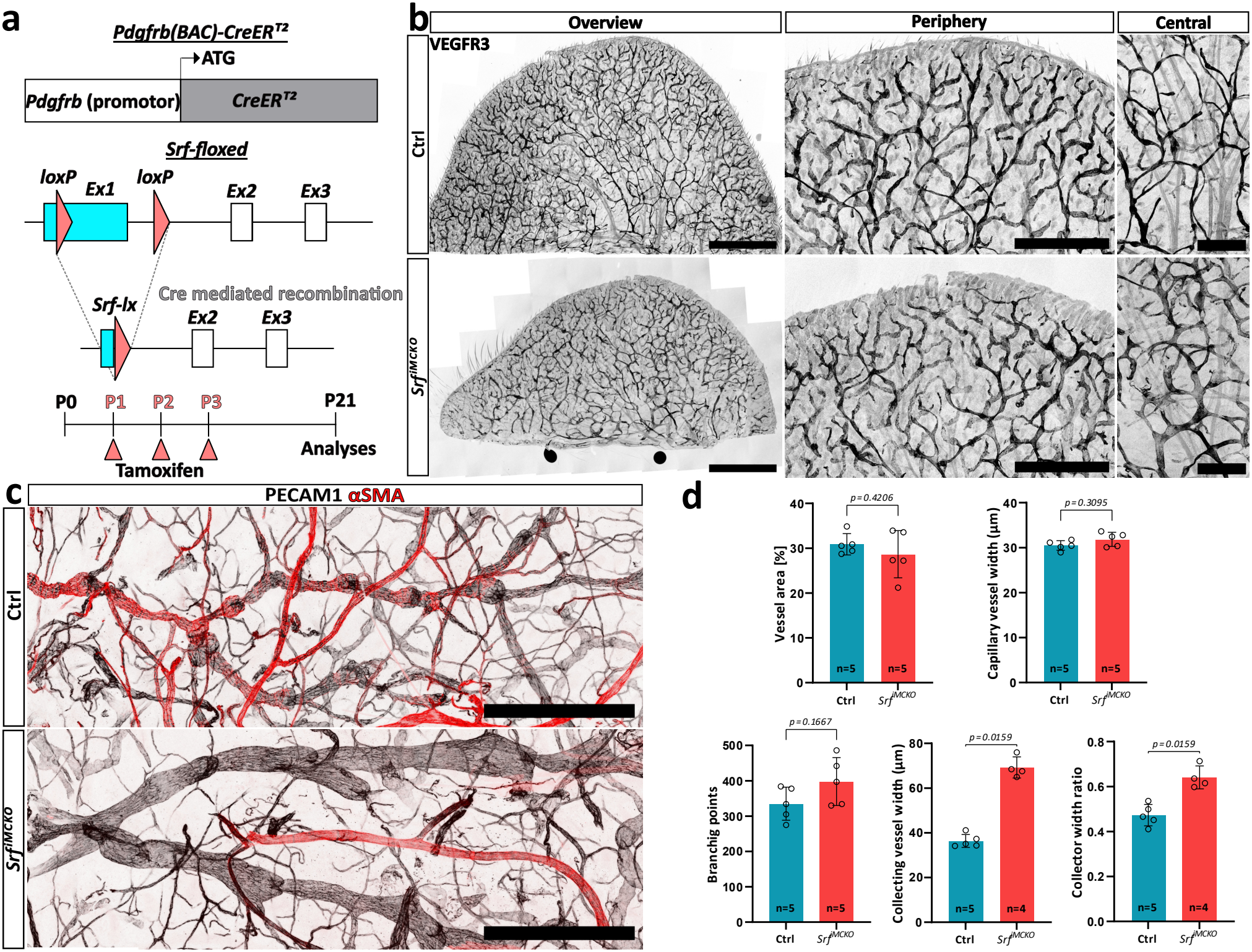
Mural cell SRF deletion selectively alters the morphology of lymphatic collecting vessels while preserving lymphatic capillaries. **(a)** Schematic of the *Pdgfrb(BAC)-CreER*^*T2*^ transgene and Cremediated recombination of the *Srf*-floxed allele (upper), and tamoxifen administration timeline (lower). **(b)** Confocal images of whole-mount ear tissues stained for VEGFR3 (gray) to visualize the lymphatic vascular plexus. Scale bars, 2 mm (left), 1 mm (middle), and 500 µm (right). **(c)** Confocal images showing lymphatic branches in control (Ctrl) and *Srf*^*iMCKO*^ ear tissues stained for PECAM1 (gray) and αSMA (red). Scale bar, 500 µm. **(d)** Morphometric analysis of vessel area (number of animals (n)= 5, each group), capillary vessel width (n=5, each group), branching points (n=5, each group), collecting vessel width (n=5 Ctrls and n=4 *Srf*^*iMCKO*^) and collecting width ratio (n=5 Ctrls and n=4 *Srf*^*iMCKO*^) of the lymphatic ear vasculature in Ctrl and Srf^iMCKO^ mice. Morphometric data showing preserved lymphatic capillary morphology but significant alterations in collecting vessel morphology. Error bars represent SD. Statistical comparison was performed using an unpaired Mann-Whitney U-test. P-values are indicated.

To characterize the lymphatic vasculature, we performed whole-mount immunofluorescence staining followed by confocal microscopy. Staining for the lymphatic endothelial marker VEGFR3 revealed no apparent differences in the lymphatic capillary network between *Srf*^*iMCKO*^ mice and littermate controls. Morphometric analysis confirmed that lymphatic capillary parameters were largely unaffected by *Srf* deletion (Figure 2b, d). In contrast, collecting vessels in *Srf*^*iMCKO*^ ear tissues exhibited striking morphological abnormalities, characterized by severe dilation and an irregular vessel shape (Figure 2b-d). High-resolution tile scans of ear tissues stained for the pan-endothelial marker PECAM1 in combination with the contractile protein αSMA revealed a loss of the characteristic segmented morphology of collecting vessels, accompanied by pronounced vessel dilation (Figure 2c). Morphometric analysis confirmed a significant increase in collecting vessel width and vessel width ratio, but no significant changes in number of branching points in *Srf*^*iMCKO*^ mice compared with littermate controls (Figure 2d).

### *Srf* ^iMCKO^ mutant lSMCs fail to express key contractile genes, exhibit abnormal morphology, and display reduced proliferative capacity

Similar to our previous observations in retinal blood vessels^30^, we observed an almost complete absence of αSMA-positive cells surrounding *Srf*^*iMCKO*^ lymphatic collecting vessels (Figure 3a). To determine whether lSMCs were absent from the collecting vessels or had instead lost αSMA expression, we extended our immunofluorescence analysis to include Desmin, a cytoskeletal protein whose expression is independent of SRF function^30^. This allowed us to identify lSMCs independently of their expression of αSMA. While Desmin staining confirmed the presence of lSMCs on *Srf*^*iMCKO*^ collecting vessels, we observed a moderate overall reduction in lSMC coverage compared with control vessels (Figure 3a, b). Additional staining for the contractile apparatus proteins Transgelin (also known as SM22α) and Calponin similarly revealed a pronounced loss of these characteristic SMC markers in *Srf*^*iMCKO*^ vessels (Figure 3c). We did not identify lSMCs that retained expression of any of the tested SRF-dependent contractile proteins, further supporting efficient *Srf* deletion in lSMCs.

**Figure 3.**
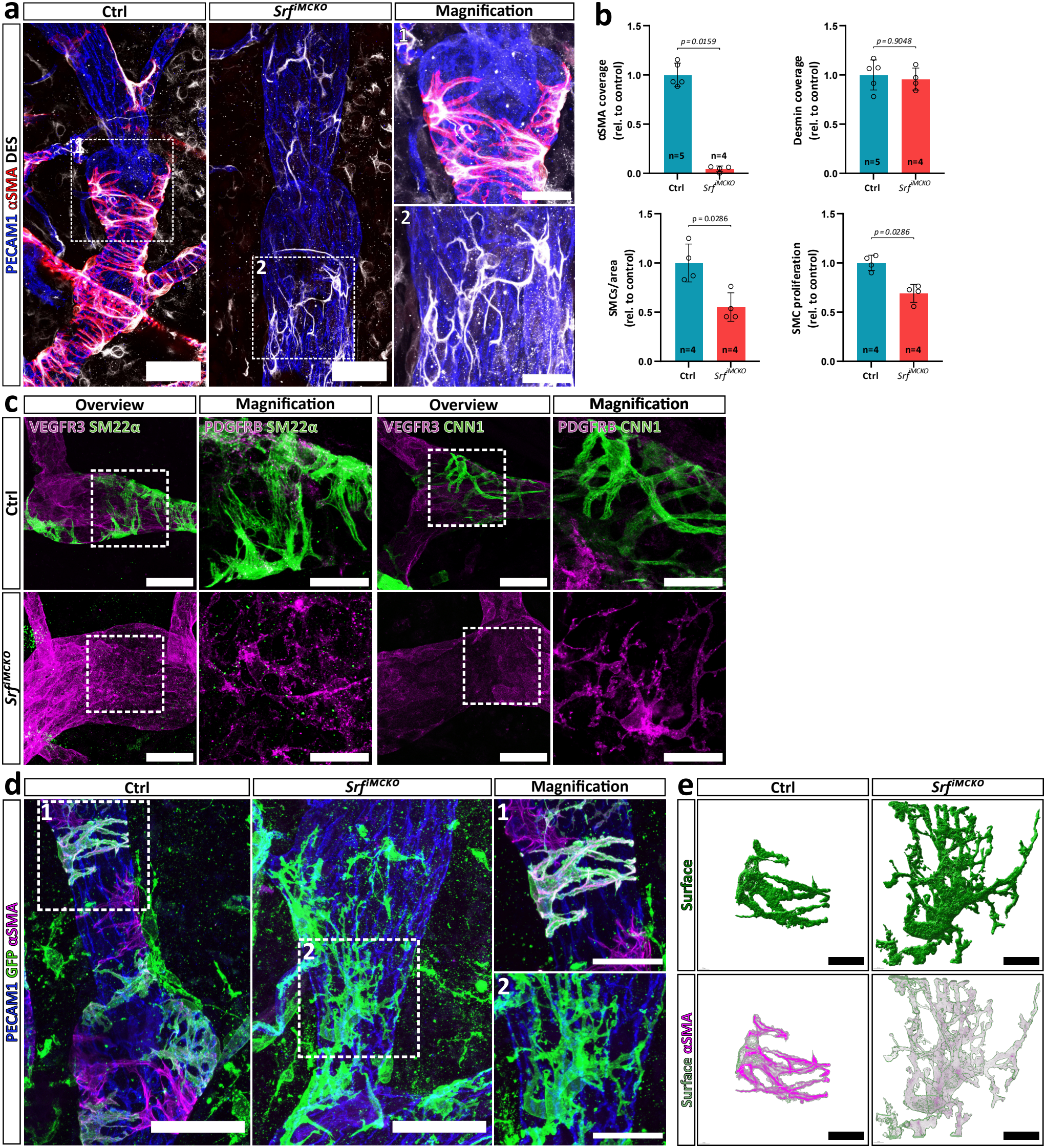
SRF regulates smooth muscle gene expression and morphology in lymphatic SMCs. **(a)** High-resolution confocal images of control (Ctrl) and *Srf*^*iMCKO*^ lymphatic collecting vessels showing lymphatic SMC coverage. Staining for PECAM1 (blue), αSMA (red), and Desmin (white). Magnified regions are outlined. Scale bars, 50 µm (main) and 25 µm (magnification). **(b)** Quantification of αSMA coverage (number of analyzed animals (n)=5 Ctrl and n=4 *Srf*^*iMCKO*^), Desmin coverage (n=5 Ctrl and n=4 *Srf*^*iMCKO*^), SMCs (n=4, each group) and SMC proliferation (n=4, each group) on lymphatic collecting vessels. **(c)** High-resolution confocal images of Ctrl and *Srf*^*iMCKO*^ lymphatic collecting vessels showing loss of characteristic SMC-markers in *Srf*^*iMCKO*^ vessels. Staining for PDGFRβ (magenta), SM22α (green), and Calponin (green). Magnified regions are outlined. Scale bars, 50 µm (main) and 25 µm (magnification). **(d)** High-resolution confocal images of lymphatic SMCs labeled with the *mTmG*-reporter (green) in collecting vessels. PECAM1 (blue) and αSMA (magenta) label endothelial cells and SMCs, respectively. Magnified regions are outlined. Scale bars, 50 µm (main) and 25 µm (magnification). **(e)** Three-dimensional (3D) renderings of representative control (Ctrl) and *Srf*^*iMCKO*^ lymphatic SMCs shown in (d). Upper panels show surface views; lower panels show transparent surface renderings with αSMA (magenta) staining to visualize the intracellular content. Scale bar, 5 µm. Error bars represent SD. Statistical comparisons were performed using an unpaired Mann-Whitney U-test. The number of analyzed animals (n) is indicated. P-values are indicated.

At higher magnification, we observed pronounced morphological alterations in *Srf*^*iMCKO*^ lSMCs (Figure 3a, magnified panels). Compared with control cells, *Srf*^*iMCKO*^ lSMCs exhibited longer and more disorganized cellular protrusions. To further characterize lSMC morphology, we took advantage of the *Rosa*^*mTmG*^ reporter allele, a genetic tool that enables ubiquitous expression of membrane-targeted tdTomato before Cre-mediated recombination and membrane-targeted enhanced green fluorescent protein (eGFP) following recombination^47^. The membrane-localized eGFP signal provides a clear outline of individual cell morphology and therefore enables detailed analysis of cellular architecture. We crossed the *Rosa*^*mTmG*^ reporter allele into the Srf-floxed background and analyzed lSMCs in *Pdgfrb-CreER*^*T2*^; *Rosa*^*mTmG*^; *Srf*^fl/fl^ (*Srf*^*iMCKO*^) mice and *Pdgfrb-CreER*^*T2*^; *Rosa*^mTmG^; *Srf*^fl/wt^ littermate controls. Control lSMCs exhibited tightly apposed, finger-like cellular protrusions extending around the collecting vessels, consistent with their characteristic organization around the vessel wall (Figure 3d). In contrast, *Srf*^*iMCKO*^ lSMCs displayed thin and elongated protrusions with an irregular and highly branched organization. Three-dimensional (3D) reconstruction further highlighted these morphological abnormalities and demonstrated the localization of αSMA within the cellular protrusions of control lSMCs and its absence in *Srf*^*iMCKO*^ mutant cells (Figure 3e).

Given the reduced expression of cytoskeletal and contractile proteins and the altered cellular morphology, we next investigated whether *Srf* deletion also affected lSMC proliferation. We administered 5-ethynyl-2⍰-deoxyuridine (EdU), a thymidine analog incorporated into newly synthesized DNA, to label proliferating cells. Combined immunostaining for VEGFR3 to identify lymphatic endothelial cells and Desmin to identify lSMCs allowed us to quantify proliferating lSMCs. We observed a statistically significant but moderate reduction of approximately 30% in lSMC proliferation in *Srf*^*iMCKO*^ collecting vessels compared to controls (Supplementary Figure 1).

Taken together, these findings demonstrate that SRF is required to maintain the expression of key contractile genes in lSMCs. Loss of SRF also results in profound alterations in lSMC morphology and a moderate reduction in lSMC proliferative capacity, indicating that SRF contributes to the maintenance of multiple cellular properties of lymphatic SMCs.

### *Srf* ^iMCKO^ lymphatic collecting vessels are enlarged but maintain lymphatic endothelial differentiation

Proper recruitment of lSMCs is crucial for the morphogenesis and maturation of lymphatic collecting vessels^6, 19, 46^. We therefore investigated whether the pronounced morphological abnormalities of *Srf*^iMCKO^ collecting vessels were associated with defects in lymphatic endothelial maturation or identity. Distinguishing between these possibilities was essential before attributing the phenotype to impaired contraction: because collecting vessel maturation itself depends on lSMC recruitment^46^, the malformations we observed could in principle arise from defective lymphatic endothelial specification rather than from the loss of lSMC contractility. The lymphatic vessel endothelial hyaluronan receptor 1 (LYVE1) is expressed throughout the lymphatic network but is downregulated specifically during the maturation of collecting vessels, while remaining expressed in lymphatic capillaries. This downregulation coincides with the recruitment of lSMCs to developing collecting vessels^46^. We therefore performed immunofluorescence staining of *Srf*^*iMCKO*^ and control ear tissues for the pan-lymphatic marker podoplanin (PDPN) in combination with LYVE1. We observed no significant differences in LYVE1 expression or distribution between the two genotypes (Figure 4a, e). In both control and *Srf*^*iMCKO*^ tissues, LYVE1 was readily detected in lymphatic capillaries, whereas collecting vessels were largely LYVE1-negative. These findings indicate that, despite the severe morphological abnormalities, *Srf*^*iMCKO*^ collecting vessels undergo the characteristic downregulation of LYVE1 associated with collecting vessel maturation.

**Figure 4.**
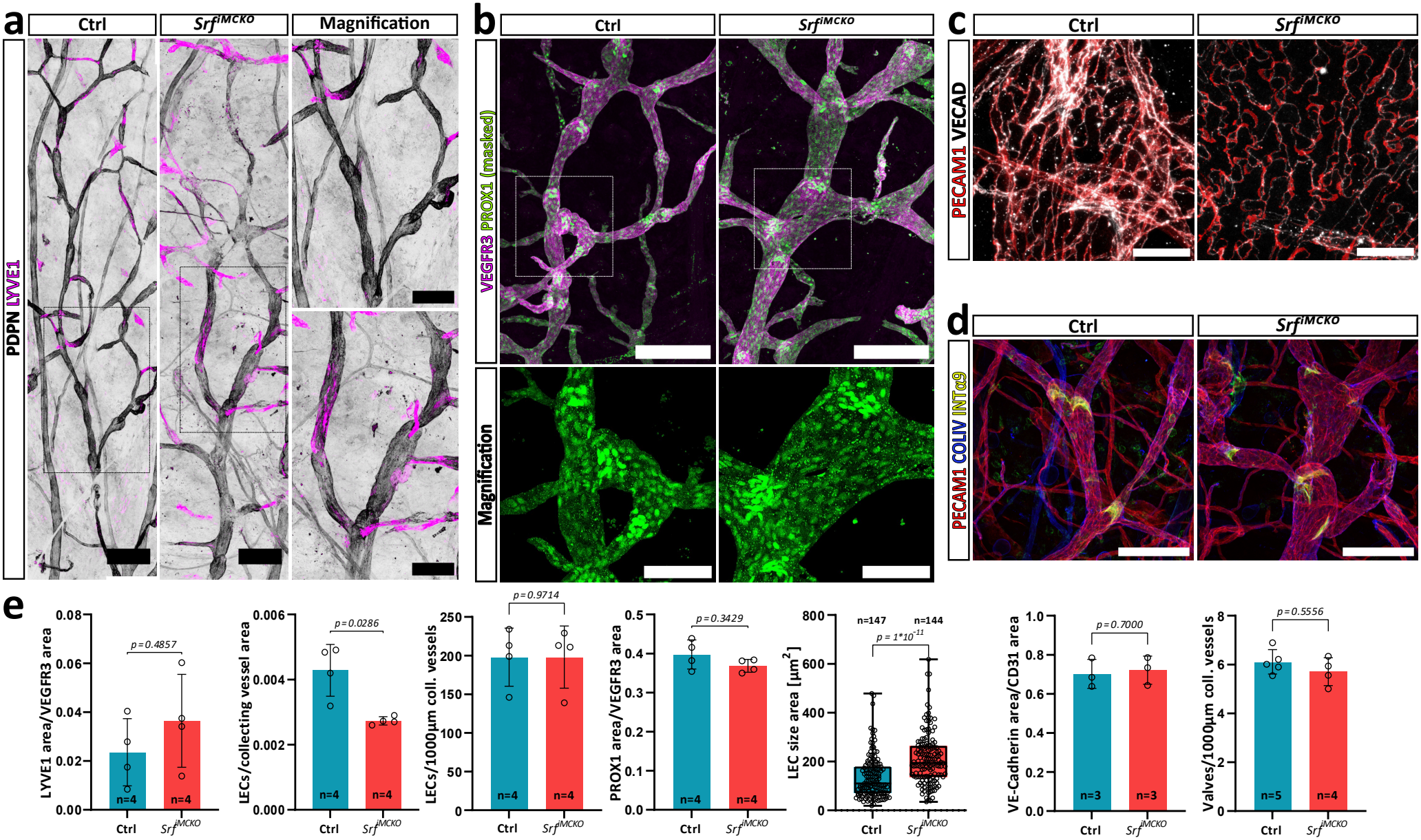
Structural characteristics of *Srf*^*iMCKO*^ lymphatic collecting vessels. **(a)** Confocal images of lymphatic branches stained for podoplanin (PDPN, gray) and LYVE1 (magenta). Magnified regions are outlined. Scale bars, 250 µm (main) and 150 µm (magnification). **(b)** Confocal overview and high-resolution images of lymphatic collecting vessels stained for VEGFR3 (magenta) and PROX1 (green). Magnified regions are outlined. Scale bars, 250 µm (main) and 100 µm (magnification). **(c)** High-resolution confocal images of endothelial junctional organization in collecting vessels stained for PECAM1 (red) and VE-Cadherin (white). Scale bar, 25 µm. **(d)** Confocal images of lymphatic valve organization stained for Integrin α9 (ITGA9, yellow), PECAM1 (red), and collagen IV (COLIV, blue). Scale bar, 100 µm. **(e)** Quantification of lymphatic vessel features. LYVE1/VEGFR3 area (number of analyzed animals (n)=4, each group), LECs/ collecting vessel area (n=4, each group), LECs/1000µm (n=4, each group), PROX1 area (n=4, each group), LEC size area (number of analyzed cells= 147 Ctrl and 144 *Srf*^*iMCKO*^, based on 3 animals each group), VE-Cadherin area (n=3, each group) and Valves/1000µm (n=5 Ctrls and n=4 *Srf*^*iMCKO*^) Error bars represent SD. Statistical comparisons were performed using an unpaired Mann-Whitney U-test. The number of analyzed animals (n) is indicated. P-values are indicated.

Lymphatic valve development is another hallmark of collecting vessel specification and is influenced by lymphatic flow^18-20^, which may be altered in *Srf*^*iMCKO*^ mutants because of their pronounced vessel abnormalities. We therefore assessed lymphatic valve formation by immunostaining for integrin α9 (ITGA9), which is highly expressed in lymphatic valves^23, 48^, together with the extracellular matrix protein collagen IV (COLIV) and PECAM1. Imaging confirmed the presence of lymphatic valves at the expected locations within collecting vessels in *Srf*^*iMCKO*^ mice (Figure 4d, e). We also found no significant differences in valve abundance between *Srf*^*iMCKO*^ and control vessels (Figure 4e). Together, these findings indicate that lymphatic endothelial specification and key features of collecting vessel maturation are largely preserved following mural cell-specific *Srf* deletion. Defective lymphatic endothelial specification is therefore unlikely to account for the collecting vessel abnormalities of these mutants, and loss of lSMC contractility itself seems the most likely explanation.

We next investigated whether the pronounced collecting vessel dilation could be explained by changes in endothelial cell number or size. We performed immunofluorescence staining for prospero Homeobox 1 (PROX1), a transcription factor required for the establishment and maintenance of lymphatic endothelial cell identity^1, 49-51^, to quantify lymphatic endothelial cells (LECs) in *Srf*^*iMCKO*^ and control collecting vessels. While the total number of LECs per collecting vessel length was unchanged, the number of LECs relative to vessel area was reduced in *Srf*^*iMCKO*^ vessels (Figure 4b, e), suggesting an increase in endothelial cell size. To further validate this finding, we stained for VE-cadherin, an endothelial adherens junction protein that delineates endothelial cell borders. Quantification of individual collecting LECs confirmed a significant increase in cell size in *Srf*^*iMCKO*^ mutants (Figure 4c, e). In addition, *Srf*^*iMCKO*^ collecting vessels displayed a partial transition from the characteristic zipper-like endothelial junctional organization of collecting vessels toward a puzzle like pattern typically observed in lymphatic capillaries^42^. This altered junctional organization may reflect changes in the mechanical environment of the enlarged collecting vessels and associated endothelial remodeling. Together with the preserved endothelial identity described above, these observations position the endothelial changes as a consequence, rather than a cause, of the collecting vessel phenotype.

### Loss of mural SRF leads to impaired lymphatic drainage

The loss of contractile proteins and the pronounced morphological abnormalities of *Srf*^*iMCKO*^ lSMCs suggested impaired contractile function, which may contribute to the severe dilation of collecting vessels. We therefore wanted to directly assess lSMC contractile function in vivo using multiphoton-guided intravital microscopy. The thin and flat anatomy of the ear skin makes it particularly well suited for this minimally invasive imaging approach.

Mice were anesthetized and fluorescently labeled dextran was injected subcutaneously into the tip of the ear. The dye is taken up by lymphatic capillaries and subsequently transported through pre-collecting and collecting vessels, allowing visualization of lymphatic drainage and collecting vessel contractions. Intravital imaging of control collecting vessels revealed slow, regular, and periodic changes in vessel diameter, consistent with rhythmic lymphatic pumping. In contrast, such periodic contractions were not detectable in *Srf*^*iMCKO*^ collecting vessels (Figure 5a, Video 1). Quantification of vessel diameter over time confirmed the absence of the characteristic diameter oscillations in *Srf*^*iMCKO*^ vessels (Figure 5a, T1 and T2; Figure 5b).

**Figure 5.**
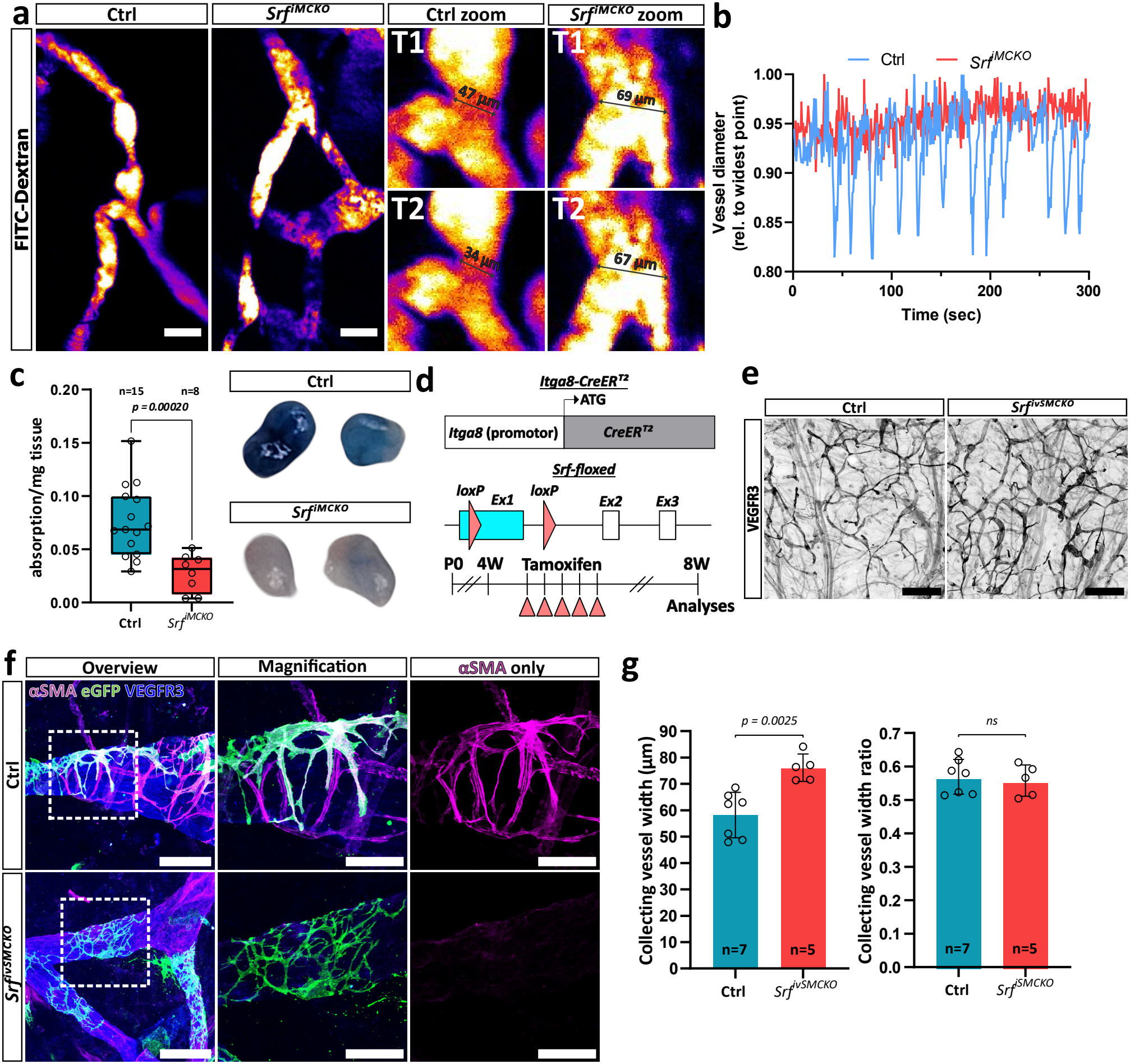
SRF regulates lymphatic vessel contraction and maintains the contractile phenotype of lymphatic SMCs. **(a)** Intravital confocal imaging of control and *Srf*^*iMCKO*^ collecting vessels following FITC-Dextran injection into the ear skin. T1 and T2 indicate representative time points during vessel contraction and relaxation, respectively. Scale bar, 100 µm. **(b)** Exemplary quantification of vessel diameter in a control and an *Srf*^*iMCKO*^ collecting vessel. **(c)** Functional assessment of lymphatic drainage following subcutaneous injection of Evans blue dye into the tail (see schematic in Figure 1b) and quantification of Evans blue content in inguinal lymph nodes (number of analyzed animals (n)= 15 Ctrls and 8 *Srf*^*iMCKO*^). Inguinal lymph nodes were collected 5 min after injection. **(d)** Schematic of the Itga8-CreER^T2^ transgene, Cre-mediated recombination of the *Srf*-floxed allele, and tamoxifen administration timeline (lower) for inducible *Srf* deletion in adult lymphatic SMCs. **(e)** Confocal images of whole-mount ear tissues stained for VEGFR3 (gray) to visualize the lymphatic vascular plexus. Scale bars, 500 µm. **(f)** High-resolution confocal images of lymphatic SMCs labeled with the mTmG reporter (green) in collecting vessels of *Srf*^*ivSMCKO*^ mice and respective controls 4 weeks after *Srf*-deletion. PECAM1 (blue) labels endothelial cells, and αSMA (magenta) labels lymphatic SMCs. (g) Morphometric analysis of collecting vessel diameter (n=7 Ctrls and 5 *Srf*^*ivSMCKO*^) and vessel width and width ratio (n=7 Ctrls and 5 *Srf*^*ivSMCKO*^) 4 weeks after *Srf*-deletion in lymphatic SMCs. Error bars represent SD. Statistical comparisons were performed using an unpaired Student’s t-test with Welch’s correction (c) and the Mann-Whitney U-test (g). The number of analyzed animals (n) is indicated. P-values are indicated.

To directly assess lSMC contractile activity at the single-cell level, we again took advantage of the *Rosa*^*mTmG*^ reporter system to visualize Cre-recombined, membrane-targeted eGFP-positive lSMCs. In combination with subcutaneous injection of fluorescently labeled dextran to visualize the collecting vessel lumen, this approach enabled high-resolution intravital imaging of individual lSMCs and their surrounding vessels (Video 2). In control vessels, we observed both periodic vessel contractions and smaller, localized contractile movements of individual lSMCs (Video 2). In contrast, *Srf*^*iMCKO*^ lSMCs displayed no detectable contractile movements, further supporting a severe impairment of lSMC contractility following *Srf* deletion (Video 2).

Although fluorescently labeled dextran could still be observed within the collecting vessels of *Srf*^*iMCKO*^ mice, indicating that lymphatic transport was not completely abolish, we next determined whether the overall efficiency of lymphatic drainage was impaired. We therefore performed the Evans blue lymphatic drainage assay described above (Figure 1b), using a longer drainage distance to more sensitively assess lymph transport. Consistent with the impaired contractile activity observed by intravital imaging, Evans blue content in the inguinal lymph nodes was significantly reduced in *Srf*^*iMCKO*^ mice compared with controls (Figure 5c). These findings demonstrate that mural cell-specific *Srf* deletion impairs lymphatic drainage and support an important contribution of lSMC-mediated vessel contraction to effective lymph transport.

### SRF function is required to maintain the contractile phenotype of lymphatic SMCs after its establishment

The distinction between establishing the contractile phenotype and maintaining it is central to the model suggested by our aging data: if declining SRF activity contributes to the age-associated loss of lSMC contractility, then SRF must be required continuously in mature lSMCs, and not only for their initial differentiation. To determine whether SRF is required not only during postnatal development but also for the maintenance of the lSMC contractile phenotype after it is established, we crossed *Srf*-floxed mice with *Itga8-CreER*^*T2*^ mice (Figure 5d). This recently published Cre line is driven by the Itga8 promoter and enables efficient recombination in vSMCs and lSMCs in adult mice, while largely sparing visceral SMCs^39^. Importantly, this approach avoids the intestinal obstruction associated with *Srf* deletion using conventional SMC-specific Cre lines, such as *Myh11-CreER*^*T2*^ or *Tagln-CreERT2* ^52, 53^.

To induce Srf deletion in vSMCs and lSMCs, 4-week-old *Itga8-CreER*^*T2*^*;Srf*^*fl/fl*^ mice and corresponding *Rosa*^*mTmG*^ reporter mice were treated with tamoxifen by five consecutive intraperitoneal injections (*Srf*^*ivSMCKO*^). Ear tissues were analyzed 4 weeks after the final tamoxifen injection (Figure 5d). Consistent with previous findings by Warthi et al. (2022) ^39^, we observed no signs of intestinal obstruction following *Srf* deletion. Immunofluorescence staining for VEGFR3 and subsequent morphometric analysis revealed a mild but significant enlargement of lymphatic collecting vessels (Figure 5e, g). Analysis of the *Rosa*^*mTmG*^ reporter mice, combined with staining for VEGFR3 and αSMA, further demonstrated a loss in αSMA expression and morphological alterations of lSMCs similar to those observed following postnatal *Srf* deletion (Figure 5f). These findings indicate that SRF is required not only for the establishment of the lSMC contractile phenotype during postnatal development but also for its maintenance thereafter.

## Discussion

Here we identify the loss of lSMC contractility, rather than the loss of lSMCs themselves, as the principal change underlying the age-associated decline of collecting vessel function, and we identify serum response factor (SRF) as the transcription factor that sustains this contractile phenotype. In aged mice, collecting vessels dilated, contractile protein expression fell and lymphatic drainage was impaired, while lSMCs identified independently of αSMA remained largely on the vessel wall; SRF expression declined in parallel. Inducible genetic deletion of *Srf* in mural cells reproduced this state, demonstrating that SRF is required for the canonical smooth muscle contractile gene programme and for normal lSMC morphology, collecting vessel organization and lymphatic pumping. Deletion of *Srf* in already-established lSMCs using *Itga8*-CreERT2 caused the same loss of αSMA expression and a similar morphological change, showing that SRF is required not only during postnatal development but also for maintenance of the contractile phenotype thereafter, and providing a model in which age-associated lymphatic decline can be studied genetically.

Our analysis of aged mice is consistent with previous reports describing reduced lSMC coverage and lymphatic pumping in aged mice and rats^12-15^. In addition, our Evans blue assay revealed reduced lymphatic drainage in aged animals, linking the structural changes to impaired lymphatic function. Previous studies have largely attributed reduced αSMA expression during aging to loss of lSMCs^12-16^. In contrast, by identifying lSMCs independently of αSMA using Desmin and integrin α8, we found that lSMC coverage of aged collecting vessels was only modestly reduced, whereas αSMA expression within the retained cells was markedly lower and increasingly heterogeneous. The simultaneous reduction in SRF expression and in established SRF-regulated contractile genes in aged lSMCs therefore provides a potential molecular mechanism underlying the age-associated decline in the lSMC contractile phenotype.

Deletion of *Srf* in mural cells resulted in a pronounced loss of αSMA, Transgelin/SM22α, and Calponin, while lSMCs remained present on collecting vessels. Persistence of the cells alongside loss of their contractile programme is the hallmark of phenotypic modulation, long described for vascular smooth muscle cells switching between contractile and synthetic states during remodeling and disease^27, 54^; our data indicate that lymphatic muscle cells are subject to an equivalent, SRF-dependent control. SRF-deficient lSMCs additionally displayed abnormal cellular morphology and moderately reduced proliferation, indicating that SRF regulates multiple aspects of lSMC homeostasis. Loss of the contractile phenotype was accompanied by severe dilation and morphological abnormalities of collecting vessels, whereas lymphatic capillaries remained largely unaffected. The simplest explanation is that a vessel that can no longer generate tone distends passively under filling pressure and remodels into the irregular, dilated shape we observe. This mirrors the retina, where loss of myogenic activity in vSMCs triggers arteriovenous shunts that relieve the resulting rise in pressure^30^: in both vascular beds, removing SRF-dependent contractility leaves the vessel unable to oppose the pressure within it, and the vessel remodels accordingly. Despite these pronounced abnormalities, *Srf*^*iMCKO*^ vessels maintained key features of lymphatic endothelial specification and maturation, including LYVE1 downregulation, valve formation, and overall LEC numbers. The increased vessel diameter was instead associated with enlarged LECs and altered endothelial junctional organization, suggesting that endothelial changes arise secondary to the altered vessel mechanics that follow the loss of lSMC contractility. Lymphatic endothelial cells actively remodel their shape and junctional organization in response to mechanical load^42^, so both the increase in cell size and the partial shift towards a capillary-like junctional pattern are consistent with a collecting vessel endothelium adapting to a wall that no longer contracts.

Our findings are consistent with previous studies demonstrating that proper mural cell recruitment is required to constrain the calibre of lymphatic collecting vessels. Loss of endothelial PDGFB, which prevents lSMC recruitment, results in similarly dilated collecting vessels^6^. That study also reported a loss of contractile function in vessels lacking lSMCs, although it detected neither a corresponding reduction in lymphatic drainage nor overt lymphedema. It is worth noting, though, that the assay used by Wang et al. involved injecting a fluorescent tracer into the ear and monitoring its clearance from that same local depot by live imaging, a qualitative assessment that detected no difference between genotypes. Our assay instead quantifies how much Evans blue reaches the inguinal lymph node within 5 min of injection into the tail, capturing transport along an extended lymphatic route in the narrow window in which active, lSMC-driven pumping is expected to be rate-limiting. The reduced drainage we observe is thus not necessarily inconsistent with the earlier study, but likely reflects the greater sensitivity of a quantitative, distal assay over a qualitative, local one. Consistent with the reduced drainage, intravital imaging directly demonstrated a severe loss of rhythmic collecting vessel contractions and of any detectable contractile movement in individual *Srf*^*iMCKO*^ lSMCs. These findings support an important role for lSMC-mediated contraction in effective lymphatic transport.

Finally, the *Itga8-CreER*^*T2*^-mediated deletion experiments, performed after the lSMC contractile phenotype was already established, demonstrate that the requirement for SRF extends beyond its developmental establishment. This later *Srf* deletion resulted in reduced αSMA expression, altered lSMC morphology, and mild collecting vessel enlargement, closely resembling aspects of the postnatal *Srf*^*iMCKO*^ phenotype. Together with the age-associated reduction in SRF expression, these findings support a model in which declining SRF activity contributes to the progressive, age-associated loss of lSMC contractile function. These findings highlight SRF-dependent contractility as an important pathway for conditions associated with impaired lymphatic pumping, including chronic lymphedema, although SRF’s essential role in other muscle types^52, 53^ means that lymphatic-SMC-specific delivery tools will need to be developed before this pathway can be targeted therapeutically in disease or aging.

Several limitations should be acknowledged. Neither Cre line used here is specific to lymphatic SMCs: *Pdgfrb-CreER*^*T2*^ targets mural cells throughout the vasculature, including pericytes and vSMCs, and *Itga8-CreER*^*T2*^ recombines in vSMCs as well as lSMCs. We can therefore not formally exclude that *Srf* deletion in other mural cell populations contributes to the lymphatic phenotype. Several observations nonetheless argue that the collecting vessel defect reflects the loss of SRF in lSMCs themselves. The phenotype is confined to lSMC-covered collecting vessels, whereas lymphatic capillaries, which are not invested by mural cells, remain unaffected. Lymphatic endothelial identity, valve formation and LEC number are preserved, so the defect is not a general failure of lymphatic development. The changes we observe in lSMCs are cell-intrinsic, affecting the expression of SRF target genes and the shape of individual cells while the cells remain on the vessel wall. Deletion using an independent Cre line after the contractile phenotype is established similarly reproduces the loss of αSMA and the change in lSMC morphology. A second limitation is that the link to aging remains correlative: SRF and its target genes decline in aged lSMCs, and removing SRF is sufficient to abolish contractility, but we have not tested whether restoring SRF activity in aged animals rescues lymphatic pumping. Finally, although lSMCs remain in place, *Srf* deletion also moderately reduces their coverage and proliferation, so contractility is not the only property that changes in these mutants. Testing whether sustaining SRF activity preserves lymphatic pumping in aged animals is the obvious next step, and would establish whether the decline we describe is a cause of age-associated lymphatic failure rather than a correlate of it. None of these caveats diminish the central implication of our findings: that SRF-dependent lSMC contractility is a distinct, and in principle modifiable, contributor to lymphatic pumping, one that acts independently of the vessel-patterning and mural-cell-recruitment defects that dominate most models of lymphatic dysfunction.

## Figure legends

**Supplementary Figure 1** | **SRF-deficient lymphatic SMCs exhibit reduced proliferation**. (a) Confocal images of proliferating lymphatic SMCs (EdU, yellow) in control (Ctrl) and *Srf*^*iMCKO*^ collecting vessels, stained for αSMA (red) or Desmin (magenta) and VEGFR3 (blue). Scale bars, 250 µm (main) and 50 µm (magnification). (b) Quantification of lymphatic SMC proliferation (number of analyzed animals (n)=4, each group). Error bars represent SD. Statistical comparison was performed using the Mann-Whitney U test. The number of analyzed animals (n) is indicated. P-values are indicated.

**Video 1** | **Rhythmic contraction of collecting lymphatic vessels is lost in** *Srf*^*iMCKO*^ mice. Multiphoton intravital imaging of an ear collecting vessel in a control and an *Srf*^*iMCKO*^ mouse following subcutaneous injection of FITC-dextran into the tip of the ear. Control vessels undergo slow, regular, periodic changes in diameter, which are not detectable in *Srf*^*iMCKO*^ vessels. Time stamp show min:sec, scale bar as indicated. Video is related to Figure 5a,b.

**Video 2** | **Individual lymphatic SMCs contract in control but not in** Srf^iMCKO^ **collecting vessels**. Multiphoton intravital imaging of *Rosa*^*mTmG*^-labelled lSMCs (green) in ear collecting vessels of a control and an *Srf*^*iMCKO*^ mouse, with the vessel lumen visualized by subcutaneously injected fluorescently labelled dextran. Control lSMCs show periodic vessel contractions together with smaller, localized contractile movements; no contractile movement is detectable in *Srf*^*iMCKO*^ lSMCs. Time stamp show min:sec, scale bar as indicated. Video is related to Figure 5.

## Notes

### Competing Interest Statement

The authors have declared no competing interest.

## References

1. Alitalo K, Tammela T, Petrova TV. Lymphangiogenesis in development and human disease. Nature 2005;438:946–953.

2. Petrova TV, Koh GY. Biological functions of lymphatic vessels. Science 2020;369.

3. Petrova TV, Koh GY. Organ-specific lymphatic vasculature: From development to pathophysiology. J Exp Med 2018;215:35–49.

4. Bachmann SB, Gsponer D, Montoya-Zegarra JA, Schneider M, Scholkmann F, Tacconi C, Noerrelykke SF, Proulx ST, Detmar M. A Distinct Role of the Autonomic Nervous System in Modulating the Function of Lymphatic Vessels under Physiological and Tumor-Draining Conditions. Cell Rep 2019;27:3305–3314 e3313.

5. Zawieja SD, Castorena-Gonzalez JA, Scallan JP, Davis MJ. Differences in L-type Ca(2+) channel activity partially underlie the regional dichotomy in pumping behavior by murine peripheral and visceral lymphatic vessels. Am J Physiol Heart Circ Physiol 2018;314:H991–H1010.

6. Wang Y, Jin Y, Mae MA, Zhang Y, Ortsater H, Betsholtz C, Makinen T, Jakobsson L. Smooth muscle cell recruitment to lymphatic vessels requires PDGFB and impacts vessel size but not identity. Development 2017;144:3590–3601.

7. Grada AA, Phillips TJ. Lymphedema: Pathophysiology and clinical manifestations. J Am Acad Dermatol 2017;77:1009–1020.

8. Arroyo-Ataz G, Jones D. Overview of Lymphatic Muscle Cells in Development, Physiology, and Disease. Microcirculation 2024;31:e12887.

9. Davis MJ, Kim HJ, Nichols CG. K(ATP) channels in lymphatic function. Am J Physiol Cell Physiol 2022;323:C1018–C1035.

10. Davis MJ, Kim HJ, Zawieja SD, Castorena-Gonzalez JA, Gui P, Li M, Saunders BT, Zinselmeyer BH, Randolph GJ, Remedi MS, Nichols CG. Kir6.1-dependent K(ATP) channels in lymphatic smooth muscle and vessel dysfunction in mice with Kir6.1 gain-of-function. J Physiol 2020;598:3107–3127.

11. Ogata F, Fujiu K, Koshima I, Nagai R, Manabe I. Phenotypic modulation of smooth muscle cells in lymphoedema. Br J Dermatol 2015;172:1286–1293.

12. Du T, Raghunandan A, Mestre H, Pla V, Liu G, Ladron-de-Guevara A, Newbold E, Tobin P, Gahn-Martinez D, Pattanayak S, Huang Q, Peng W, Nedergaard M, Kelley DH. Restoration of cervical lymphatic vessel function in aging rescues cerebrospinal fluid drainage. Nat Aging 2024;4:1418–1431.

13. Kataru RP, Park HJ, Shin J, Baik JE, Sarker A, Brown S, Mehrara BJ. Structural and Functional Changes in Aged Skin Lymphatic Vessels. Front Aging 2022;3:864860.

14. Akl TJ, Nagai T, Cote GL, Gashev AA. Mesenteric lymph flow in adult and aged rats. Am J Physiol Heart Circ Physiol 2011;301:H1828–1840.

15. Lei PJ, Ruscic KJ, Roh K, Rajotte JJ, O’Melia MJ, Bouta EM, Marquez M, Pereira ER, Kumar AS, Razavi MS, Zhou H, Menzel L, Huang L, Kumra H, Duquette M, Huang P, Baish JW, Munn LL, Kurpios NA, Ubellacker JM, Padera TP. Aging-induced changes in lymphatic muscle cell transcriptomes are associated with reduced pumping of peripheral collecting lymphatic vessels in mice. Dev Cell 2024.

16. Nagai T, Bridenbaugh EA, Gashev AA. Aging-associated alterations in contractility of rat mesenteric lymphatic vessels. Microcirculation 2011;18:463–473.

17. Gonzalez-Loyola A, Bovay E, Kim J, Lozano TW, Sabine A, Renevey F, Arroz-Madeira S, Rapin A, Wypych TP, Rota G, Durot S, Velin D, Marsland B, Guarda G, Delorenzi M, Zamboni N, Luther SA, Petrova TV. FOXC2 controls adult lymphatic endothelial specialization, function, and gut lymphatic barrier preventing multiorgan failure. Sci Adv 2021;7.

18. Sabine A, Agalarov Y, Maby-El Hajjami H, Jaquet M, Hagerling R, Pollmann C, Bebber D, Pfenniger A, Miura N, Dormond O, Calmes JM, Adams RH, Makinen T, Kiefer F, Kwak BR, Petrova TV. Mechanotransduction, PROX1, and FOXC2 cooperate to control connexin37 and calcineurin during lymphatic-valve formation. Dev Cell 2012;22:430–445.

19. Sabine A, Bovay E, Demir CS, Kimura W, Jaquet M, Agalarov Y, Zangger N, Scallan JP, Graber W, Gulpinar E, Kwak BR, Makinen T, Martinez-Corral I, Ortega S, Delorenzi M, Kiefer F, Davis MJ, Djonov V, Miura N, Petrova TV. FOXC2 and fluid shear stress stabilize postnatal lymphatic vasculature. J Clin Invest 2015;125:3861–3877.

20. Saygili Demir C, Sabine A, Gong M, Dormond O, Petrova TV. Mechanosensitive mTORC1 signaling maintains lymphatic valves. J Cell Biol 2023;222.

21. Frye M, Taddei A, Dierkes C, Martinez-Corral I, Fielden M, Ortsater H, Kazenwadel J, Calado DP, Ostergaard P, Salminen M, He L, Harvey NL, Kiefer F, Makinen T. Matrix stiffness controls lymphatic vessel formation through regulation of a GATA2-dependent transcriptional program. Nat Commun 2018;9:1511.

22. Sweet DT, Jimenez JM, Chang J, Hess PR, Mericko-Ishizuka P, Fu J, Xia L, Davies PF, Kahn ML. Lymph flow regulates collecting lymphatic vessel maturation in vivo. J Clin Invest 2015;125:2995–3007.

23. Bazigou E, Lyons OT, Smith A, Venn GE, Cope C, Brown NA, Makinen T. Genes regulating lymphangiogenesis control venous valve formation and maintenance in mice. J Clin Invest 2011;121:2984–2992.

24. Wang Z, Wang DZ, Pipes GC, Olson EN. Myocardin is a master regulator of smooth muscle gene expression. Proc Natl Acad Sci U S A 2003;100:7129–7134.

25. Sun Q, Chen G, Streb JW, Long X, Yang Y, Stoeckert CJ, Jr., Miano JM. Defining the mammalian CArGome. Genome Res 2006;16:197–207.

26. Olson EN, Nordheim A. Linking actin dynamics and gene transcription to drive cellular motile functions. Nat Rev Mol Cell Biol 2010;11:353–365.

27. Owens GK, Kumar MS, Wamhoff BR. Molecular regulation of vascular smooth muscle cell differentiation in development and disease. Physiol Rev 2004;84:767–801.

28. Miano JM. Serum response factor: toggling between disparate programs of gene expression. J Mol Cell Cardiol 2003;35:577–593.

29. Yoshida T, Owens GK. Molecular determinants of vascular smooth muscle cell diversity. Circ Res 2005;96:280–291.

30. Orlich MM, Dieguez-Hurtado R, Muehlfriedel R, Sothilingam V, Wolburg H, Oender CE, Woelffing P, Betsholtz C, Gaengel K, Seeliger M, Adams RH, Nordheim A. Mural Cell SRF Controls Pericyte Migration, Vessel Patterning and Blood Flow. Circ Res 2022;131:308–327.

31. Arroyo-Ataz G, Yague AC, Breda JC, Mazzilli SA, Jones D. Single-Cell Transcriptomics and Lineage Tracing Unveil Parallels in Lymphatic Muscle and Venous Smooth Muscle Development, Identity, and Function. Arterioscler Thromb Vasc Biol 2025;45:1207–1225.

32. Rashbrook VS, Brash JT, Ruhrberg C. Cre toxicity in mouse models of cardiovascular physiology and disease. Nat Cardiovasc Res 2022;1:806–816.

33. Schindelin J, Arganda-Carreras I, Frise E, Kaynig V, Longair M, Pietzsch T, Preibisch S, Rueden C, Saalfeld S, Schmid B, Tinevez JY, White DJ, Hartenstein V, Eliceiri K, Tomancak P, Cardona A. Fiji: an open-source platform for biological-image analysis. Nat Methods 2012;9:676–682.

34. Honkura N, Richards M, Lavina B, Sainz-Jaspeado M, Betsholtz C, Claesson-Welsh L. Intravital imaging-based analysis tools for vessel identification and assessment of concurrent dynamic vascular events. Nat Commun 2018;9:2746.

35. Orlich MM, Norren A, Desai M, Holm A, Kozakewich H, Schoofs H, Makinen T, Bischoff J, Gaengel K. Intravital Imaging of Disease Mechanisms in a Mouse Model of CCM Skin Lesions-Brief Report. Arterioscler Thromb Vasc Biol 2025;45:113–118.

36. Corliss BA, Doty RW, Mathews C, Yates PA, Zhang T, Peirce SM. REAVER: A program for improved analysis of high-resolution vascular network images. Microcirculation 2020;27:e12618.

37. McDowell KP, Berthiaume AA, Tieu T, Hartmann DA, Shih AY. VasoMetrics: unbiased spatiotemporal analysis of microvascular diameter in multi-photon imaging applications. Quant Imaging Med Surg 2021;11:969–982.

38. Hao Y, Hao S, Andersen-Nissen E, Mauck WM, 3rd, Zheng S, Butler A, Lee MJ, Wilk AJ, Darby C, Zager M, Hoffman P, Stoeckius M, Papalexi E, Mimitou EP, Jain J, Srivastava A, Stuart T, Fleming LM, Yeung B, Rogers AJ, McElrath JM, Blish CA, Gottardo R, Smibert P, Satija R. Integrated analysis of multimodal single-cell data. Cell 2021;184:3573–3587 e3529.

39. Warthi G, Faulkner JL, Doja J, Ghanam AR, Gao P, Yang AC, Slivano OJ, Barris CT, Kress TC, Zawieja SD, Griffin SH, Xie X, Ashworth A, Christie CK, Bryant WB, Kumar A, Davis MJ, Long X, Gan L, Belin de Chantemele EJ, Lyu Q, Miano JM. Generation and Comparative Analysis of an Itga8-CreER (T2) Mouse with Preferential Activity in Vascular Smooth Muscle Cells. Nat Cardiovasc Res 2022;1:1084–1100.

40. Davis MJ, Kim HJ, Li M, Zawieja SD. A vascular smooth muscle-specific integrin-alpha8 Cre mouse for lymphatic contraction studies that allows male-female comparisons and avoids visceral myopathy. Front Physiol 2022;13:1060146.

41. Zawieja SD, Pea GA, Broyhill SE, Patro A, Bromert KH, Norton CE, Kim HJ, Sivasankaran SK, Li M, Castorena-Gonzalez JA, Drumm BT, Davis MJ. Cellular characterization of the mouse collecting lymphatic vessels reveals that lymphatic muscle cells are the innate pacemaker cells. Elife 2025;12.

42. Schoofs H, Daubel N, Schnabellehner S, Gronloh MLB, Palacios Martinez S, Halme A, Marks AM, Jeansson M, Barcos S, Brakebusch C, Benedito R, Engelhardt B, Vestweber D, Gaengel K, Linsenmeier F, Schurmann S, Saharinen P, van Buul JD, Friedrich O, Smith RS, Majda M, Makinen T. Dynamic cytoskeletal regulation of cell shape supports resilience of lymphatic endothelium. Nature 2025.

43. Schoofs H, Makinen T. Genetic targeting of lymphatic endothelial cells in mice: current strategies and future perspectives. Int J Dev Biol 2024;68:189–198.

44. Kraft M, Schoofs H, Petkova M, Andrade J, Grosso AR, Benedito R, De Roo AK, Boon LM, Vikkula M, Kapp FG, Hagerling R, Potente M, Makinen T. Angiopoietin-TIE2 feedforward circuit promotes PIK3CA-driven venous malformations. Nat Cardiovasc Res 2025;4:801–820.

45. Petkova M, Kraft M, Stritt S, Martinez-Corral I, Ortsater H, Vanlandewijck M, Jakic B, Baselga E, Castillo SD, Graupera M, Betsholtz C, Makinen T. Immune-interacting lymphatic endothelial subtype at capillary terminals drives lymphatic malformation. J Exp Med 2023;220.

46. Lutter S, Xie S, Tatin F, Makinen T. Smooth muscle-endothelial cell communication activates Reelin signaling and regulates lymphatic vessel formation. J Cell Biol 2012;197:837–849.

47. Muzumdar MD, Tasic B, Miyamichi K, Li L, Luo L. A global double-fluorescent Cre reporter mouse. Genesis 2007;45:593–605.

48. Bazigou E, Xie S, Chen C, Weston A, Miura N, Sorokin L, Adams R, Muro AF, Sheppard D, Makinen T. Integrin-alpha9 is required for fibronectin matrix assembly during lymphatic valve morphogenesis. Dev Cell 2009;17:175–186.

49. Adams RH, Alitalo K. Molecular regulation of angiogenesis and lymphangiogenesis. Nat Rev Mol Cell Biol 2007;8:464–478.

50. Lupu IE, Grainger DE, Kirschnick N, Weischer S, Zhao E, Martinez-Corral I, Schoofs H, Vanhollebeke M, Jones G, Godwin J, Forrow A, Lahmann I, Riley PR, Zobel T, Alitalo K, Makinen T, Kiefer F, Stone OA. Direct specification of lymphatic endothelium from mesenchymal progenitors. Nat Cardiovasc Res 2025;4:45–63.

51. Tammela T, Alitalo K. Lymphangiogenesis: Molecular mechanisms and future promise. Cell 2010;140:460–476.

52. Mericskay M, Blanc J, Tritsch E, Moriez R, Aubert P, Neunlist M, Feil R, Li Z. Inducible mouse model of chronic intestinal pseudo-obstruction by smooth muscle-specific inactivation of the SRF gene. Gastroenterology 2007;133:1960–1970.

53. Angstenberger M, Wegener JW, Pichler BJ, Judenhofer MS, Feil S, Alberti S, Feil R, Nordheim A. Severe intestinal obstruction on induced smooth muscle-specific ablation of the transcription factor SRF in adult mice. Gastroenterology 2007;133:1948–1959.

54. Miano JM, Fisher EA, Majesky MW. Fate and State of Vascular Smooth Muscle Cells in Atherosclerosis. Circulation 2021;143:2110–2116.

