## Supplementary Figure 1 for "Loss of SRF-driven contractility, not lymphatic smooth muscle cell loss, is the primary driver of age-associated collecting vessel decline"

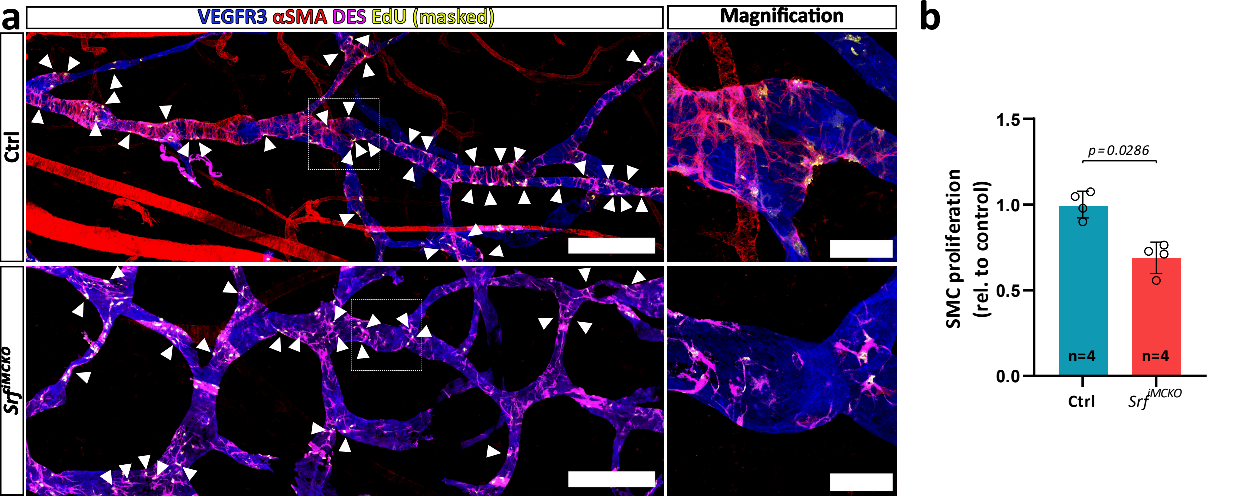
a
VEGFR3 αSMA DES EdU (masked)
Magnification
Ctrl
Srf^iMCKO
b
SMC proliferation (rel. to control)
1.5
1.0
0.5
0.0
p = 0.0286
n=4
n=4
Ctrl
Srf^iMCKO
